# Performance Evaluation of Prediction on Molecular Graphs with Graph Neural Networks

**DOI:** 10.1101/2022.10.21.513175

**Authors:** Haotian Li

## Abstract

Machine learning and deep learning are novel and trending approaches to solving real-world scientific problems. Graph machine learning is dedicated to performing learning methods, such as graph neural networks, on non-Euclidean data such as graphs. Molecules, with their natural graph structures, could be analyzed by such method. In this work, we carry out the performance evaluation regarding to learning results as well as time consumed, speedup, and efficiency using different types of neural network structures and distributed training pipeline implementations. Besides, the reasons lead to an unideal performance enhancement is investigated. Code availability at https://github.com/htlee6/perf-analysis-dist-training-gnn.

## 1 Introduction

Recent years have seen rapid development and success in the area of machine learning and deep learning. Machine learning and deep learning techniques have been widely applied in object detection, medical image analysis, natural language processing and understanding, audio recognition, recommendation systems, knowledge graphs, autonomous driving[1, 2, 3, 4, 5, 6, 7], etc. However, the sequentiality or the spacial order of the data limits the generalizability of these powerful models, for example, a convolutional neural network model cannot ingest a piece of non-Euclidean data. As a result, the problems of machine learning/deep learning on non-Euclidean data came into being, and the concept of graph machine learning, aiming at addressing this issue as a new approach, has been introduced.

In graph machine learning, the model is required to have the character of permutation invariance and equivariance. There are mainly 3 levels of tasks in the area of graph machine learning, i.e. node-level tasks, edge-level tasks, and graph-level tasks. Node-level tasks are concerned with predicting the identity or role of each node within a graph. An example of edge-level inference is predicting if an edge exists in the given graph, for example, when a new user joined Instagram, will the existing users follow the new user or vice versa? In a graph-level task, our goal is to predict the property of an entire graph. For example, for a molecule represented as a graph, we might want to predict whether it will bind to a receptor implicated in a disease.

In this work, we build a distributed training framework in order to investigate the effect of distributed training on the training performance (esp. time consumed), which is implemented using PyTorch[8] and PyTorch Ignite[9], and compare the performance difference between different graph machine learning libraries (PyG[10] and DGL[11]). From the results, we conclude that:

- Training a GIN model takes longer time than a GCN model, adding a virtual node in GNN model makes model more complex and therefore increases the training time.
- In our experiments, training with PyG is faster comparing to that using DGL with PyTorch as backend.
- The implementation of distributed training system did not achieve the expected performance, there are two known reasons: limited network communication rate and frequent switching between processes slowing the training down.

## 2 Learning on Molecular Graphs with Graph Neural Network

In this section, we are going to introduce some concepts, terminologies and models about molecular graph and graph neural network, as well as some popular libraries to build up a graph neural network model with ease.

### 2.1 Graph Representation and Molecular Graphs

Graph representation is often used in representing a graph object abstractly. A graph *G* = (*V, E*), where *V* = {1, …, *n*} is the set of vertices, and *E* ⊆ *V* × *V* is the set of edges. One way to represent a graph *G* specifically is using an adjacency matrix *A* ∈ *R*^|*V* |×|*V* |^, for example, *A*_*i,j*_ = 1 means vertex *i* and *j* are connected in an undirected graph. Another way is to use an adjacency list *L* where the elements of the list are pairs of neighbor vertices, for example, *L* = {(*i, j*), (*j, i*), (*p, q*), (*i, p*)} represents a directed graph *G* as figure 1 shows.

**Figure 1:**
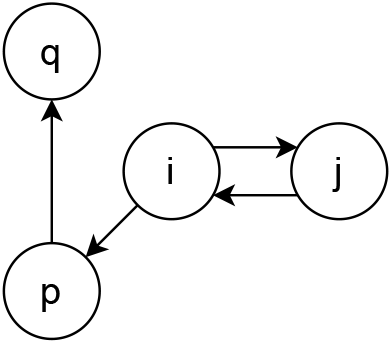
Example graph with 4 nodes, *i, j, p*, and *q*

Molecular graphs are graphs that depict the topological structures of molecules. Chemically, the vertices of a molecular graph are the elements, such as C, H, O, etc., and the edges are chemical bonds that connect the elements. Besides the adjacency relationship as information contained in the graph, the vertices and edges could also contain special information in the given context, for example, the edges/chemical bonds of a molecular graph varies depending on the elements it is connected to, the edges of a knowledge graph varies depending on the given a tri-tuple *< s, r, d >*. Node features **X**^(0)^ ∈ ℝ^|*V* |×*f*^ and edge features **E**^(0)^ ∈ ℝ^|*E*|×*d*^ could be defined with regard to the domain knowledge, where *f* and *d* indicate the dimension of each feature.

### 2.2 Graph Neural Network

A graph neural network, namely GNN, is a learning module designed for learning on graph-structured data. A GNN iteratively updates node embeddings (eq. 2) by aggregating localized information (eq. 1) via the parameterized functions.

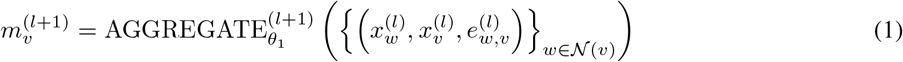

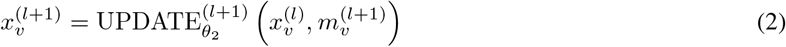

where {…} denotes a multi-set and 𝒩 (*ν*) *V* defines the neighborhood set of node *v ∈ V*. [12] After *L* times of aggregation, a graph representation could be derived via global aggregation of *X*^(*L*)^, such as summation, Set2set[13], or readout methods like virtual node[14] and etc. Figure 2 [15] illustrates the corresponding computational graph generated when calculating the embedding of target node *A* in a 2-layer GNN module.

**Figure 2:**
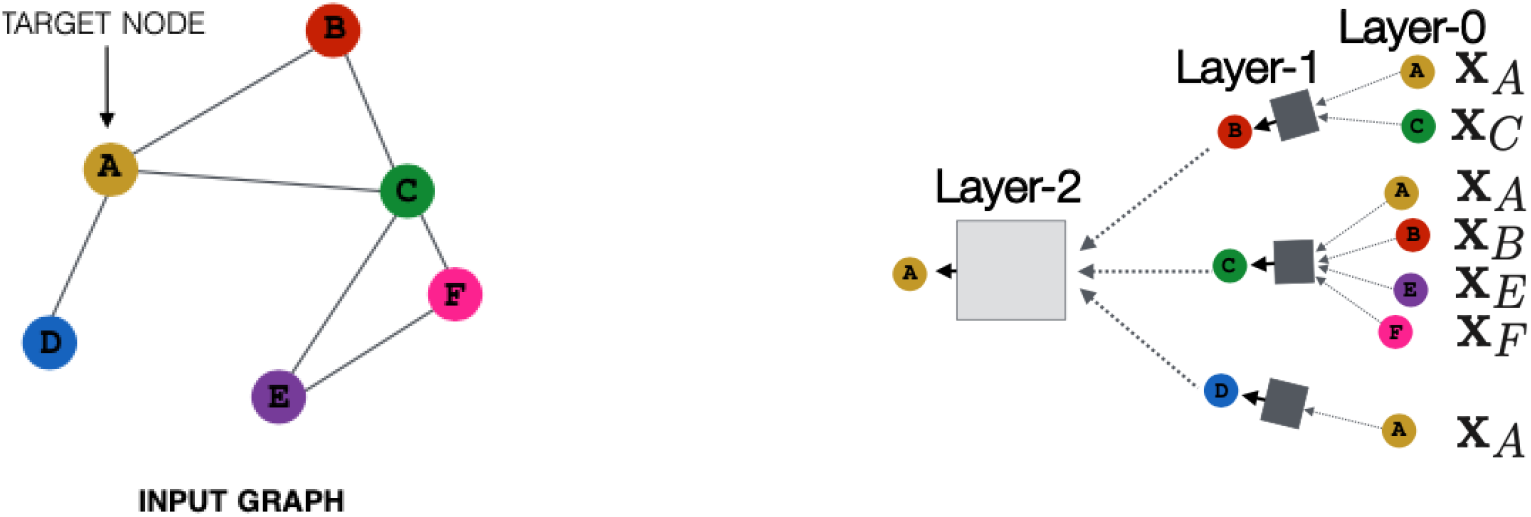
Abstract graph and corresponding computational graph

**GCN** (Graph Convolutional Network)[16] is a semi-supervised learning approach for graph-structured data. It is based on an efficient convolutional neural network variation that works directly on graphs. A localized first-order approximation of spectral graph convolutions motivates the choice of convolutional architecture. The model learns hidden layer representations that encode both local graph structure and node attributes and scales linearly in the number of graph edges.

To give a mathematical formulation of GCN along with the previously defined norm, here we can use eq. 3 and eq. 4

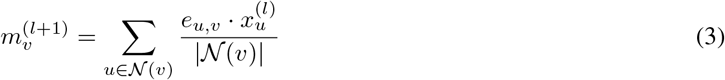

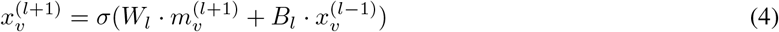

where 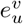 denotes the edge feature, *σ*(·) is a non-linear function, and *W*_*l*_, *B*_*l*_ are trainable weight matrix parameters. GCN could be formalized using pure matrix operations as well (eq. 5):

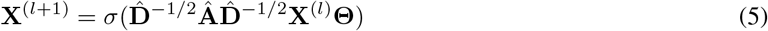

where **Â** = **A** + **I** denotes the adjacency matrix with processed self loops and 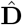 is degree matrix of the graph, which makes 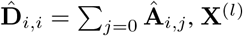 is the representation of nodes in the graph, and **Θ** denotes weight matrix.

**GIN** (Graph Isomorphism Network) [17] is proved to be able to significantly accurate to distinguish graph structures and is as powerful as the Weisfeiler-Lehman test. The node-wise formulation of GIN could written as eq. 6.

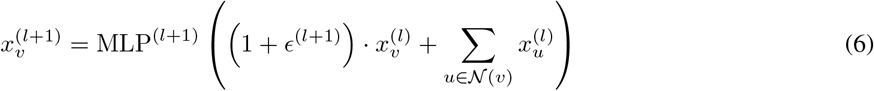

GINE convolution operator [18], as shown in eq. 7, is a variant of GIN, which is able to corporate edge features into message passing framework, and that makes GIN have a higher performance.

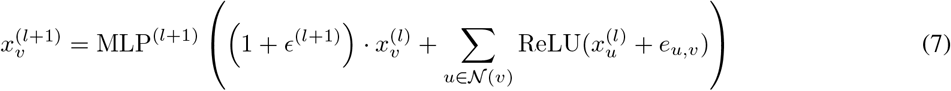

here 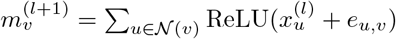 and *ϵ*^(*l*+1)^ is a trainable scalar parameter.

**Virtual Node** Different from common approaches to extract the graph-level information using simple summing up or average[19], virtual node[12] provides a novel idea to profile the embedding of the graph, which is to append a virtual node to the given homogeneous graph that is connected to all other nodes, as shown in figure 3. The virtual node serves as a global scratch space that each node both reads from and writes to in every step of message passing. This allows information to travel long distances during the propagation phase. Node and edge features of the virtual node are added as zero-filled input features. Furthermore, special edge types will be added both for in-coming and out-going information to and from the virtual node.

**Figure 3:**
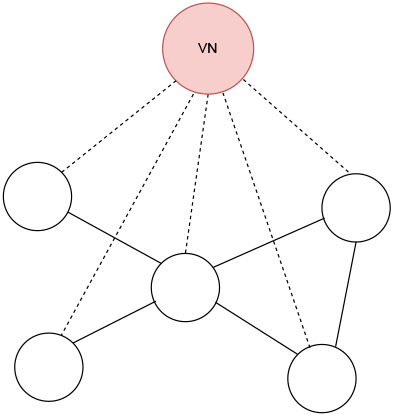
Illustration: Adding a virtual node in a 5-node graph

### 2.3 Graph Neural Network Libraries

**PyG** (PyTorch Geometric) [10] is a library built upon PyTorch to easily compose and build up GNN learning pipelines for a wide range of applications related to structured data. It consists of various methods for deep learning on graphs and other irregular structures, also known as geometric deep learning, from a variety of published papers. In addition, it consists of easy-to-use mini-batch loaders for operating on many small and single giant graphs, multi GPU-support, DataPipe support, distributed graph learning via Quiver, a large number of common benchmark datasets (based on simple interfaces to create your own), the GraphGym experiment manager, and helpful transforms, both for learning on arbitrary graphs as well as on 3D meshes or point clouds.

**DGL** (Deep Graph Library)[11] is a Python package built for easy implementation of graph neural network model family, on top of existing DL frameworks (currently supporting PyTorch, MXNet and TensorFlow). It offers a versatile control of message passing, speed optimization via auto-batching and highly tuned sparse matrix kernels, and multi-GPU/CPU training to scale to graphs of hundreds of millions of nodes and edges[20].

## 3 Distributing Training and Systems

### 3.1 Distributed Training

Training deep learning models takes time. Deep neural networks often consist of millions or billions of parameters that are trained over huge datasets. As deep learning models become more complex, computation time can become unwieldy. Training a model on a single GPU can take weeks. Distributed training can split up the training workload and share among multiple hosts (computation nodes) that could work in parallel to fix this problem.

There are two main types of distributed training: data parallelism and model parallelism. In data parallelism, the dataset is divided into partitions, which will be distributed to every node later. The model is copied in each of the nodes, and is trained on the subset of the data it received. Each node independently computes the loss and gradients and then communicate with each other to keep their corresponding model parameters, or gradients synchronized at the end of the batch computation. In model parallelism, the model is segmented into several parts that can run sequentially in different nodes, and each one of them will run on the same data. Generally, it is more difficult to implement model parallelism than data parallelism.

In distributed training, speedup and efficiency are intuitive metrics measuring the relative performance improvement as the scale of computation resource enlarges. Mathematically speaking, speedup *S*(*x*) = *T* (1)*/T* (*x*) and efficiency *E*(*x*) = *S*(*x*)*/x*, where *T* (*x*) is time consumed using *x* unit(s) of computation resource.

### 3.2 Distributed Training using PyTorch

PyTorch Distributed Data-Parallel Training (DDP) is a widely adopted single-program multiple-data training paradigm. With DDP, the model is replicated on every process, and every model replica will be fed with a different set of input data samples. DDP takes care of gradient communication to keep model replicas synchronized and overlaps it with the gradient computations to speed up training. During the backwards pass, gradients from each node are averaged.

**PyTorch Ignite** [9] is a high-level library to help with training and evaluating neural networks written in PyTorch. Besides, it also provides high-level wrappers for PyTorch multiprocessing and DDP workflows.

## 4 Dataset

**ogbg-molhiv** is a public dataset offered by Open Graph Benchmark (OGB)[21], adapted from MoleculeNet. MoleculeNet[22] is a benchmark specially designed for testing machine learning methods of molecular properties. As the authors aim to facilitate the development of molecular machine learning method, this work curates a number of dataset collections, creates a suite of software that implements many known featurizations and previously proposed algorithms.

The HIV dataset was introduced by the Drug Therapeutics Program (DTP) AIDS Antiviral Screen, which tested the ability to inhibit HIV replication for over 40,000 compounds. Screening results were evaluated and placed into three categories: confirmed inactive (CI),confirmed active (CA) and confirmed moderately active (CM). The latter two labels are further combined, making it a classification task between inactive (CI) and active (CA and CM).

## 5 Experiments

### 5.1 Model Design

Our model main body consists of 3 parts (figure 4):

- ***Data transformation and featurization*** Simplified molecular-input line-entry system, known as “SMILES”, is a specification in the form of a line notation for describing the structure of chemical species using short ASCII string, which was first initiated by David Weininger[23] in the 1980s. The data transformation and featurization of nodes and edges are processed with RDKit[24] to graph-structured objects.
- ***GNN convolutional layer stack*** The GNN convolutional layer stack consists of sequential GNN convolutional layers, batch norm layers[25], dropout layers[26] and residual connections[27]. The whole stack serves as a way to iterate and abstract the embedding of the nodes in the graph for downstream tasks.
- ***Pooling layer and linear layer*** Pooling layer profiles the graph using node embeddings and then pass the output to the linear layer, which outputs the finalized prediction of the task.

**Figure 4:**
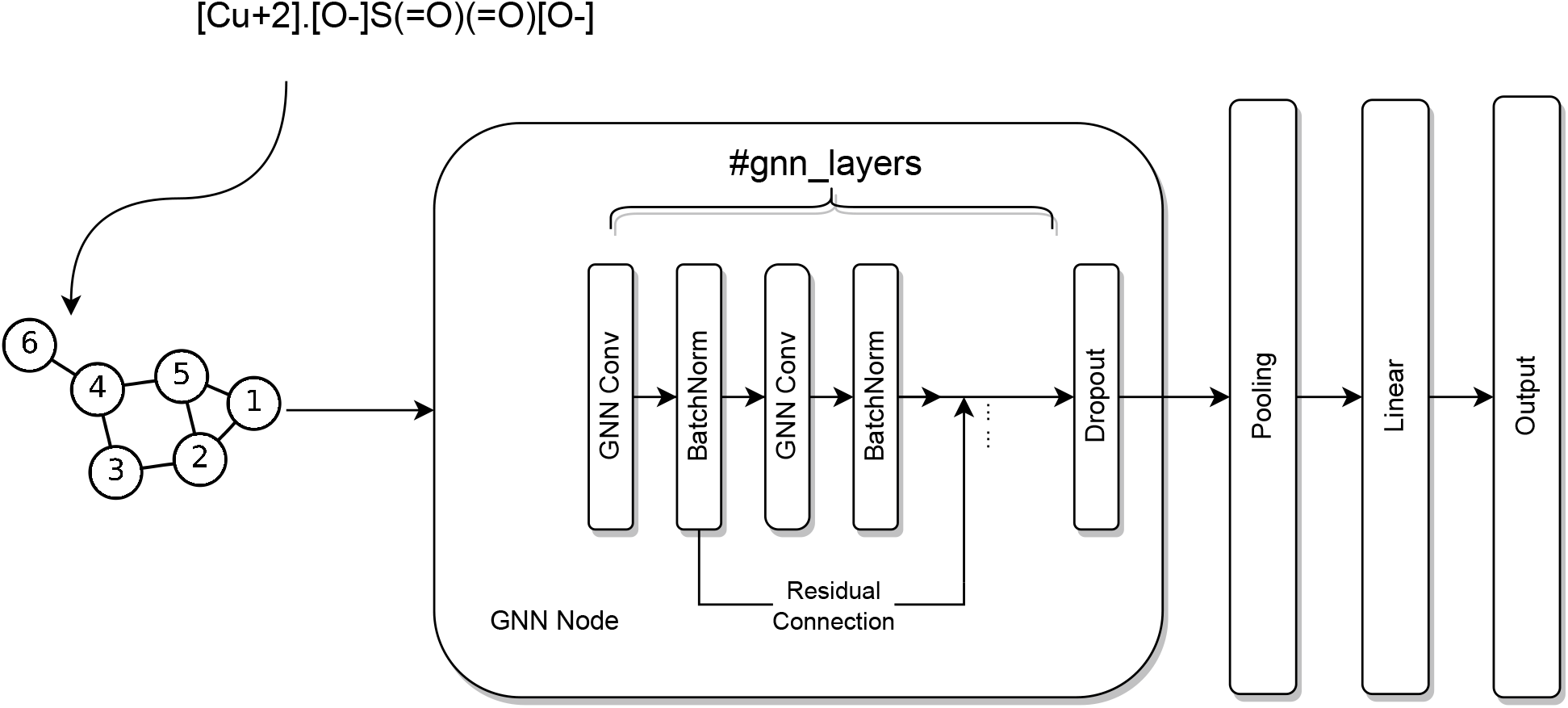
Architecture of designed model

### 5.2 Experiment Setup and Hyperparameters

We primarily conduct experiments in the modes of: (I) PyG (II) DGL with PyTorch backend (III) PyG distributed mode (using Ignite) on a cluster with the scale of 1 (standalone and multiprocessing), 2, or 3.

To build up a distributed cluster, we use 3 computing instances of ‘lrz.medium’ size on LRZ (Leibniz-Rechenzentrum), and each of them has 2 VCPUs, 9GB RAM, 20GB disk space with Ubuntu-18.04-LTS-bionic installed. The instances within the cluster communicated with each other via Munich Research Network (MWN). The hyperparameters in the learning process include: batch size=32, embedding dim=300, GNN layers=5, learning rate=0.001, dropout ratio=0.5, unless additionally specified.

### 5.3 Results

To investigate the performances in objective training modes, we implemented training modes mentioned above (Sec. 5.2) and plot the average time consumed per epoch (figure 5) in these modes and the evaluation of speedup and efficiency (figure 6) in parallel computing scenes. Training program implemented using PyG is generally faster than using DGL with PyTorch backend. PyG implementation is 16%, 11%, and 4% faster than DGL one when using GCN, GCN virtual node, and GIN model respectively. Training a GCN model is always faster than training a GIN model, time is 34%, 18%, and 60% longer when training on PyG locally, DGL and PyG distributedly. Adding virtual node to GNN model increases the training time. Table 1 shows four models could achieve similar performance result with regard to AUROC metric on test data, but GIN outperforms than others.

**Figure 5:**
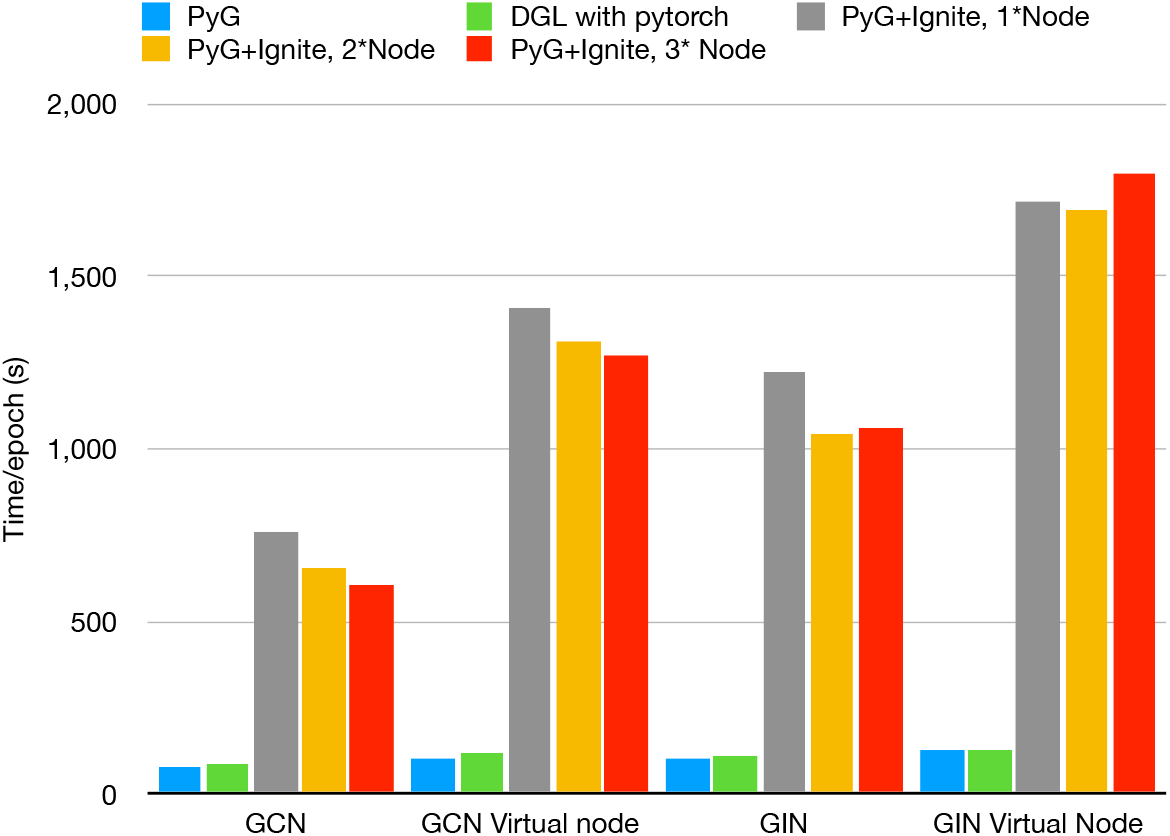
Training time in different training modes

**Figure 6:**
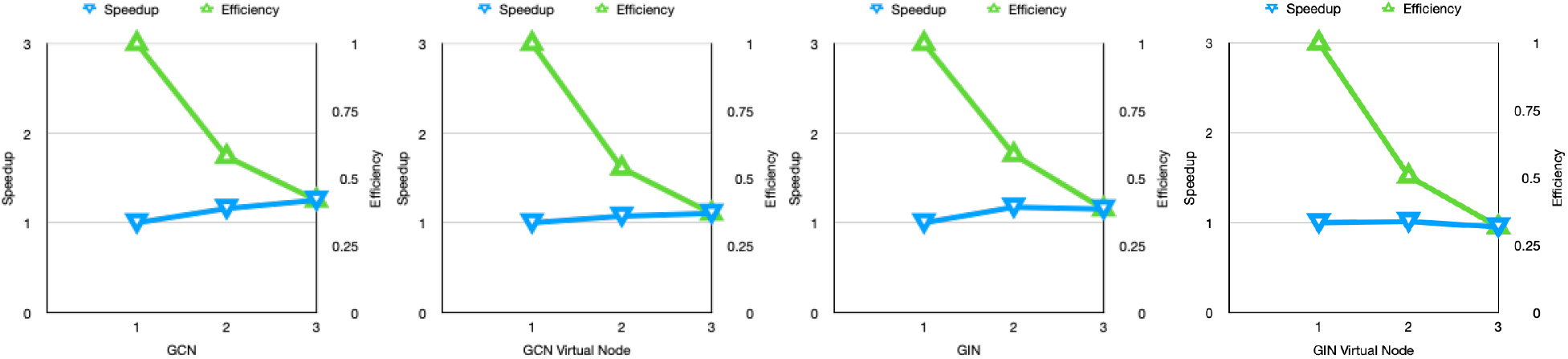
Speedup and efficiency plot in different training modes

**Table 1:** Model prediction performance

|  | GCN | GCN VN | GIN | GIN VN |
| --- | --- | --- | --- | --- |
| AUROC | 0.767 | 0.764 | 0.787 | 0.769 |

It can also be obviously observed that using distributed training increases the training time significantly. In figure 5, the training time in a 3-node cluster using PyG+Ignite takes nearly 7.8 times of that using PyG locally, and in figure 6, the increase of speedup is minimal as we increase the number of nodes within the cluster, for example, the GCN model only has a speedup of *S*(2) =1.15 and *S*(3)=1.25 which is against our intention of design. According to the discuss in PyTorch community[28], there are 3 time-consuming phases in a PyTorch distributed training loop, i.e. model forward, loss backward, and optimizer step. One possible and usual reason is that the lower rate of network communication slows down the overall training progress. Another possible reason is that frequently switching among the processes will be a heavy burden for the CPUs to work, and therefore slows down the training process.

To address this problem, we further record and analyze the time consumed in these phases, as table 2 shows. Comparing PyG, PyG+Ignite Local, PyG+Ignite Standalone Single processing, we can observe that using Ignite locally almost has no effect on training in all phases, however, PyG+Ignite Standalone Single processing mode slightly increases the training time and takes less time in forward but more time in loss backward. Comparing PyG+Ignite Standalone Single processing and PyG+Ignite Standalone Multiprocessing, it takes in all phases noticeably more time in multiprocessing mode, which explains process switching in multiprocessing mode consumes plenty of time. Comparing PyG+Ignite Standalone Multiprocessing and PyG+Ignite Distributed with 3 nodes, the rise in both absolute time from 201.84s to 387.15s and relative proportion of overall time from 34.51% to 64.49% of backward phase while the time of forward and optimizer step decreases could be explained by the communication delay caused by network when the data parallel strategy is adopted, since all communication in PyTorch DDP happens in backward phase and the rate of network becomes the bottleneck.

**Table 2:** Average time consumed of one iteration (batch size=32) in model forward, loss backward, and optimizer step phases in different settings.

|  | PyG |  | PyG+Ignite Local |  |
| --- | --- | --- | --- | --- |
|  | Time (s) | Percent | Time (s) | Percent |
| Forward | 42.14 | 54.71% | 40.31 | 54.68% |
| Backward | 29.69 | 38.55% | 28.79 | 39.05% |
| Optim Step | 5.18 | 6.72% | 4.61 | 6.25% |
| Sum | 77.01 |  | 73.72 |  |

|  | PyG+Ignite Standalone<br>Single processing |  | PyG+Ignite Standalone<br>Multiprocessing |  | PyG+Ignite<br>Distributed, 3 Nodes |  |
| --- | --- | --- | --- | --- | --- | --- |
|  | Time (s) | Percent | Time/process (s) | Percent | Time/process (s) | Percent |
| Forward | 24.89 | 30.01% | 303.96 | 51.98% | 183.37 | 30.54% |
| Backward | 49.80 | 60.03% | 201.84 | 34.51% | 387.15 | 64.49% |
| Optim Step | 8.25 | 9.94 % | 78.92 | 13.49% | 29.80 | 4.96% |
| Sum | 82.95 |  | 584.72 |  | 600.32 |  |

## 6 Conclusion

In this work, we implemented GNN (distributed) model training pipelines using PyG, DGL, and PyTorch Ignite, and evaluated the training performance among different libraries and distributing systems. Our conclusions include GIN is more powerful to discriminate graphs, using PyG can make model training faster, network conditions have a crucial effect on distributed training system, and multiprocessing can slow down the training process.

## Acknowledgments

This report is submitted in partial fulfillment of the assignment required by the Advantage Analytics and Machine Learning 2022SS course at Ludwig-Maximilians-Univeristät München.

## Notes

### Competing Interest Statement

The authors have declared no competing interest.

### Summary of Updates

Author information update.

